# Fate of the M-phase-assembled centrioles during the cell cycle in the *TP53;PCNT;CEP215*-deleted cells

**DOI:** 10.1101/2020.09.14.297440

**Authors:** Gee In Jung, Kunsoo Rhee

## Abstract

Cancer cells frequently include supernumerary centrioles. Here, we generated *TP53;PCNT;CEP215* triple knockout cell lines and observed precocious separation and amplification of the centrioles at M phase. Many of the triple KO cells maintained supernumerary centrioles throughout the cell cycle. The M-phase-assembled centrioles lack an ability to function as templates for centriole assembly during S phase. They also lack an ability to organize microtubules in interphase. However, we found that a fraction of them acquired an ability to organize microtubules during M phase. Our works provide an example how supernumerary centrioles behave in dividing cells.

## INTRODUCTION

Centrosome is a subcellular organelle which functions as a major microtubule organizing center. Centrosome is comprised of a pair of centrioles surrounded by pericentriolar material (PCM). Centriole duplication and separation are controlled in close link to the cell cycle. When a cell enters S phase, daughter centrioles start to assemble in a perpendicular angle to mother centrioles and elongate. Entering mitosis, the daughter centriole disengages from the mother centriole but remains associated until the end of mitosis (Cabral et al., 2013; Shukla et al., 2015; Seo et al., 2015). Centriole separation initiates at anaphase when separase cleaves pericentrin (PCNT), a major PCM protein (Lee and Rhee, 2012; Matsuo et al., 2012). Cleavage of PCNT induces disintegration of mitotic PCM, resulting in release of a daughter centriole from the mother centriole during mitotic exit (Kim et al., 2015).

When a cell exits M phase, a daughter centriole becomes a young mother centriole. Centriole-to-centrosome conversion is a process for a centriole to acquire PCM for microtubule organization activity (Wang et al., 2011). In addition, a young mother centriole can function as a template for procentriole assembly at subsequent S phase (Wang et al., 2011). One of the highlight events in the conversion may be accumulation of CEP152, an adaptor for PLK4 into the young mother centriole (Wang et al., 2011; Novak et al., 2014). CEP152 is recruited after a sequential process which begins immediately after daughter centriole assembly at S phase (Fu et al., 2016; Tsuchiya et al., 2016). Cep63 and Cep152 form a stable complex at the proximal end of the centrioles, and Cep57 is a proximity interactor of Cep63 (Firat-Karalar and Stearns, 2014; Kim et al., 2019; Zhao et al., 2020).

PLK4 is a central regulator of centriole duplication and, therefore, the levels and activity of PLK4 are tightly regulated during the cell cycle (O’Connell et al., 2001; Bettencourt-Diaz et al., 2005; Habedanck et al., 2005). Centriole assembly cannot start at G1 phase since the PLK4 activity is maintained low. Once the PLK4 activity is induced near S phase, daughter centrioles assemble using the mother centrioles as templates. Upon binding to STIL, PLK4 is activated through trans-autophosphorylation and phosphorylates STIL, triggering the centriolar recruitment of SAS6 and cartwheel formation (Ohta et al., 2014; Dzhindshev et al., 2014; Moyer et al., 2015; Lopes et al., 2015; Arquint et al, 2015). Mother centrioles are not allowed to form new daughter centrioles at G2 and M phase, as far as they are associated with daughter centrioles in vicinity (Wong and Stearns, 2003; Kim et al., 2016).

Supernumerary centrioles are frequently observed in diverse cancer cells (Chan, 2011; Marteil et al., 2018). Several ways to generate centriole amplification have been suggested, including centriole over duplication, de novo centrosome formation, fragmentation of overly elongated centrioles and cytokinesis failure (Sabat-Pospiech et al., 2019). Most of all, overexpression of PLK4 generates multiple daughter centriole during S phase (Habedanck et al., 2005; Kleylein-Sohn et al., 2007). In fact, genetic variants near the *PLK4* gene are closely associated with aneuploidy (McCoy et al., 2015) and overexpression of PLK4 has been reported in a variety of tumor cells (Liao et al., 2019). Involvement of PLK1 on centriole amplification was also reported. PLK1 is a novel regulator of centriole elongation in human cells (Kong et al., 2014) and augmented PLK1 activity generates mitotic centriole over-elongation (Kong et al., 2020). Over-elongated centrioles can also contribute to centrosome amplification through fragmentation or the formation of multiple procentrioles along their elongated walls (Kohlmaier et al., 2009; Marteil et al., 2018).

PCM is organized as a toroid around mature centrioles in interphase, and expands into a more amorphous structure in preparation for mitosis (Fu and Glover, 2012; Lawo et al., 2012; Mennella et al., 2012). PCNT and CEP215, two major PCM proteins in the human centrosome (Doxsey et al., 1994; Dictenberg et al., 2002; Andersen et al., 2003; Fong et al., 2007), specifically interact to each other in both interphase and mitotic centrosomes and act as scaffolds for the γ-tubulin ring complex (Kim and Rhee, 2014). *CEP215* mutations cause autosomal recessive primary microcephaly (Bond et al., 2005). Likewise, *PCNT* mutations cause microcephalic osteodysplastic primordial dwarfism as well as other disease, such as cancers and mental disorders (Rauch et al., 2008; Delaval and Doxsey, 2010).

We previously reported that centrioles prematurely separate and eventually amplify when *PCNT* is deleted (Kim et al., 2019). We interpret that absence of PCNT generate defects in mitotic PCM which holds the mother and daughter centrioles together (Kim et al., 2019). When PCM is dissolved with a prolonged treatment of mitotic drugs such as STLC and nocodazole, mother centrioles are distanced from daughter centrioles and generate nascent centrioles even at M phase (Seo et al., 2015). In this work, we generated the *TP53, PCNT* and *CEP215* triple knockout (KO) cells and determined centriole amplification. The results showed that the centrioles precociously separate and amplify during M phase. We analyzed the fate of the M-phase-assembled centrioles during the cell cycle.

## RESULTS

### Generation of the *TP53;PCNT;CEP215-*deleted cells

We previously observed precocious separation and amplification of centrioles at M phase in HeLa cells with *TP53* and *PCNT* double deletions (Kim et al., 2019). To expand our understanding on PCM regulation of centriole separation and assembly, we generated the *TP53, PCNT* and *CEP215* triple KO cells, using the CRISPR/Cas9 method (Figure1 Supplement Figure 1). During the selection step, we realized that the triple KO cells failed to form a stable cell line, due to a low proliferation activity and cell apoptosis. Therefore, the triple KO cells were generated in the presence of the ectopic *PCNT* gene with a destabilization domain (*DD-PCNT*), whose expression is induced by doxycycline and shield1 (Kim et al., 2019). The ectopic *DD-PCNT* gene was hardly expressed as far as doxycycline and shield1 were absent. In fact, immunoblot analysis revealed that PCNT and CEP215 were below detection levels in the triple KO cells (Figure 1A). Immunostaining analysis also revealed that the PCNT and CEP215 signals were absent at the centrosomes of the triple KO cells (Figure 1B-D). These results indicate that the *TP53, PCNT* and *CEP215* triple KO cell lines were properly generated. Therefore, we used the triple KO line to analyze combined roles of PCNT and CEP215 for regulation of centriole behavior during mitosis.

**Figure 1.**
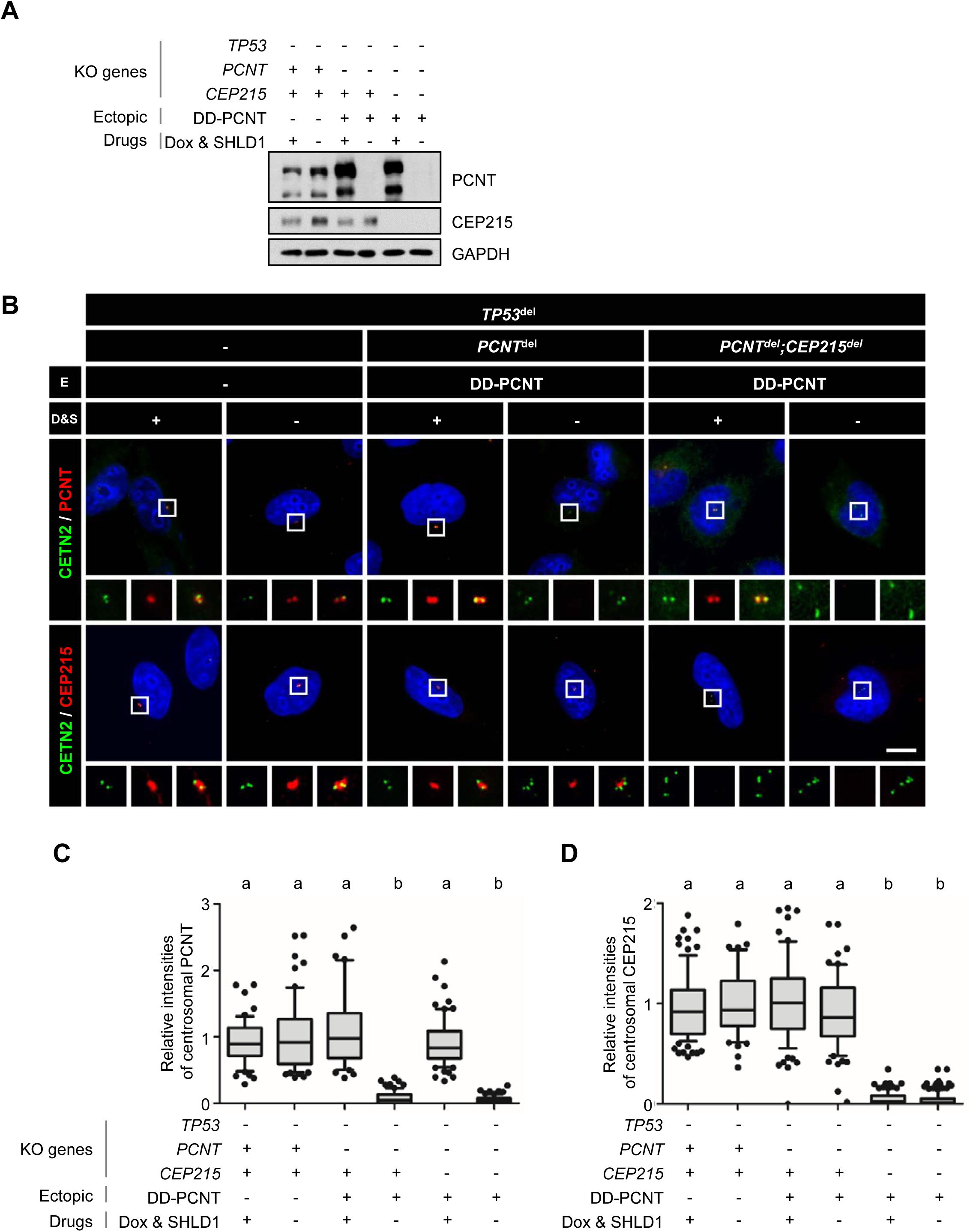
Generation of *TP53;PCNT;CEP215-*deleted cells. (A) The *TP53, PCNT* and *CEP215* genes were deleted in HeLa cells using the CRISPR/CAS9 method. Endogenous *PCNT* was deleted in the presence of ectopic *DD-PCNT* gene whose expression was induced by doxycycline (Dox) and shield1 (SHLD1). The deletions were confirmed with the immunoblot analysis with antibodies specific to PCNT, CEP215 and GAPDH. (B) The triple KO cells were coimmunostained with antibodies specific to centrin-2 (CETN2, green), PCNT (red) and CEP215 (red). Nuclei were stained with DAPI (blue). Scale bar, 10 μm. (C, D) Relative intensities of the centrosomal PCNT (C) and CEP215 (D) signals were determined. Greater than 30 cells per group were analyzed in three independent experiments. Values are means and SEM. The statistical significance was analyzed using two-way ANOVA and indicated by lower cases (P<0.05).

### Precocious centriole separation and amplification in the triple KO cells during M phase

We initially examined precocious separation and amplification of centrioles at M phase in the *TP53, PCNT* and *CEP215* triple KO cells. The cells were arrested at prometaphase using S-trityl-L-cysteine (STLC), an EG5 inhibitor, and determined centriole behaviors (Figure 2A). We confirmed that PCNT and CEP215 were not detected in the deletion cells arrested at prometaphase (Figure 2B). Precocious centriole separation was observed in most of the *TP53* and *PCNT* double KO cells and supernumerary centrioles were detected in about 40% of them (Figure 2C-E). In the *TP53* and *CEP215* double KO cells, precocious centriole separation was observed in less than a half of the cells and no centriole amplification occurred (Figure. 2C-E). Precocious centriole separation was obvious in the *TP53, PCNT* and *CEP215* triple KO cells and centriole amplification was fortified (Figure 2C-E). Indeed, some of the triple KO cells included exceeding number of extra centrioles up to 30 (Figure 2C, E). These results revealed a cooperative function of PCNT and CEP215 in prevention of precocious centriole separation and amplification during mitosis.

**Figure 2.**
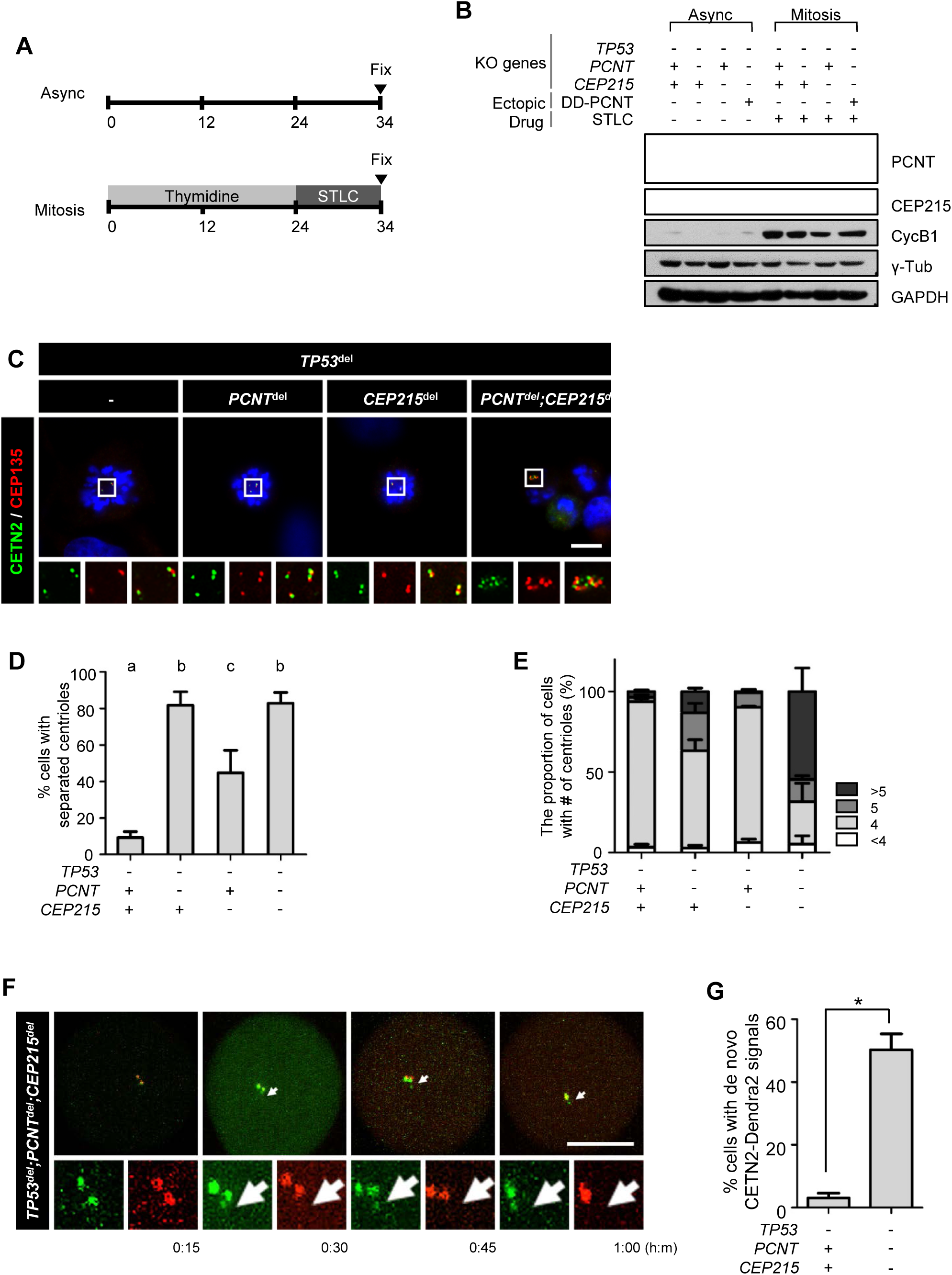
Precocious centriole separation and amplification at M phase in the triple KO cells. (A) Timeline for preparation of prometaphase cells. The triple KO cells were treated with thymidine 24 h followed by STLC for 10 h. (B) Immunoblot analyses were performed with antibodies specific to PCNT, CEP215, cyclin B1, γ-tubulin and GAPDH. (C) The prometaphase-arrested cells were subjected to coimmunostaining analysis with antibodies specific to centrin-2 (CETN2, green) and CEP135 (red). Nuclei were stained with DAPI (blue). Scale bar, 10 μm. (D) The number of cells with separated centrioles were counted, based on 1:1 ratio of the centriolar CEP135 and CETN2 signals. (E) The number of centrioles per cell were counted. Greater than 30 cells per group were analyzed in three independent experiments. Values are means and SEM. The statistical significance was analyzed using two-way ANOVA and indicated by lower cases (P<0.05). (F) The CETN2-Dendra2-expressing triple KO cells were treated with thymidine 24 h followed by STLC for 8 h. After a light activation, the CETN2-Dendra2 signals were recorded for up to 2 h. Light activation makes CETN2-Dendra2 detected with 594 nm fluorescence (red). Nascent centrioles were detected only with 488 nm fluorescence (green, white arrow). Scale bar, 10 μm. (G) The number of cells with nascent centrioles were counted. Greater than 20 cells per group were analyzed in three independent experiments. Values are means and SEM. The statistical significance was analyzed using one-way ANOVA. *, P<0.05.

We recorded generation of M-phase centrioles in the triple KO cells. We used centrin-2 coupled to the photo-convertible fluorescent protein Dendra2 (CETN2-Dendra2), which allows to distinguish newly formed centrioles from pre-existing ones (Loffler et al., 2013). The CETN2-Dendra2-expressing cells synchronously entered mitosis and were arrested at prometaphase with the STLC treatment. Nascent centrioles started to appear in prometaphase-arrested triple KO cells (Figure 2F; Figure 2 Supplement Movie 1a). However, no centriole was generated in the *TP53*-deleted control cells (Figure 2G). As far as we know, this is the first live observation of centriole generation at M phase.

### Limited centriole assembly at S phase in the triple KO cells

We traced the fate of the extra centrioles in the triple KO cells throughout the cell cycle. It is well-known that the PLK4-overexpressing cells generate extra centrioles (Habedanck et al., 2005; Coelho et al., 2015). The extra centrioles were observed when expression of the ectopic *PLK4* gene was induced by doxycycline for 24 h (Figure 3A, B). We designated the extra centrioles in the PLK4-overexpressing cells as S-phase-assembled centrioles. In comparison, the extra centrioles in the triple KO cells was designated as the M-phase-assembled centrioles.

**Figure 3.**
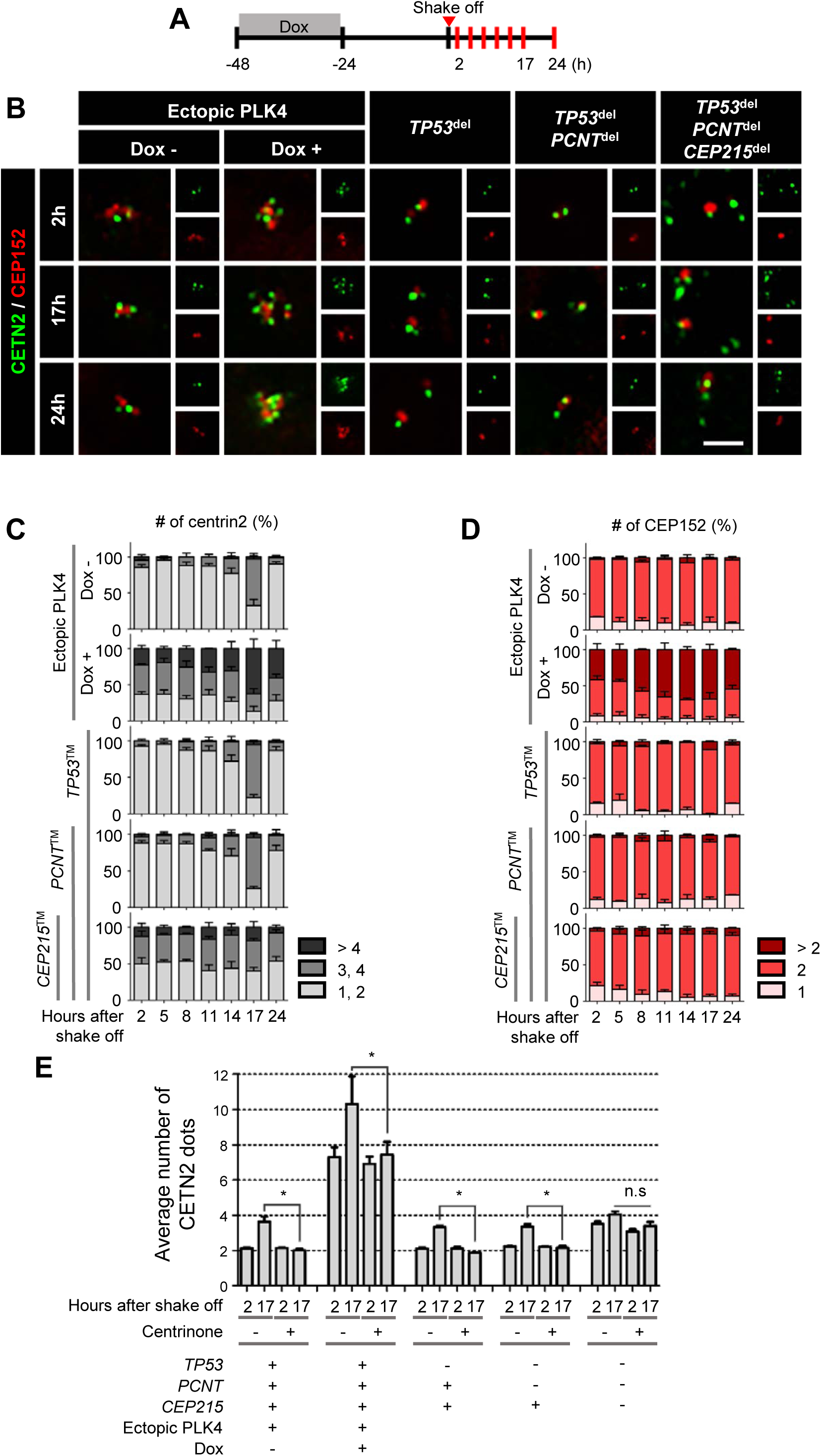
Limited S-phase centriole assembly in the triple KO cells. (A) Timeline for preparation of synchronous interphase cells. Doxycycline was treated for 24 h to induce ectopic PLK4 expression, washed out and cultured for another 24 h. Mitotic cells were collected and cultured for up to 24 h. At indicated time points, the cells were subjected to coimmunostaining analyses. (B) The PLK4-overexpressing and triple KO cells were subjected to coimmunostaining analysis with antibodies specific to CETN2 (green) and CEP152 (red). Scale bar, 2 μm. (C, D) Number of CETN2 (C) and CEP152 (D) dots were counted in the PLK4-overexpressing and triple KO cells at indicated time points. (E) After mitotic shake-off, the PLK4-overexpressing and triple KO cells were cultured in the presence of centrinone for 2 and 17 h, and immunostained with the centrin-2 antibody. The number of centrioles per cell was counted. Greater than 30 cells per group were analyzed in three independent experiments. Values are means and SEM. The statistical significance was analyzed using one-way ANOVA. *, P<0.05.

Using the mitotic shake-off method, we collected the M-phase population of the PLK4-overexpressing cells and the triple KO cells, forced them to synchronously enter into G1 phase and cultured them for up to indicated time points (Figure 3A). In control cells, the number of centrioles were two in the beginning of the culture and increased to four in 17th hour when the cells entered S phase (Figure 3B). The centriole number eventually down to two in 24th hour after the next mitosis (Figure 3B, C). The PLK4-overexpressing cells includes extra centrioles in the beginning of the culture and 60% of them had five or more centrioles at 17th hour, indicating that most of the centrioles assembled new procentrioles during the S phase (Figure 3B, C). The triple KO cells also included multiple centrioles at G1 phase. However, the number of centrioles in the triple deletion cells slightly increased at S phase (Figure 3B, C). These results suggest that the M-phase-assembled centrioles survive but do not duplicate during interphase.

CEP152 is a mother centriole protein which functions as an adaptor for PLK4 (Hatch et al., 2010). In control cells, the number of the CEP152-positive centrioles are two throughout the cell cycle (Figure 3B, D). About 40% of the PLK4-overexpressing cells included three or more CEP152-positive centrioles in the beginning of the culture and this proportion was maintained throughout the cell cycle (Figure 3B, D). This observation is consistent with the previous reports in which multiple centrioles in the PLK4-overexpressing cells can duplicate during S phase (Coelho et al., 2015). It is interesting that the triple KO cells included only two CEP152-positive centrioles out of multiple centrioles and this number was maintained throughout the cell cycle (Figure 3B, D). These results suggest that only a single out of multiple daughter centrioles can convert into a mother centriole in the triple KO cells during mitotic exit.

We used centrinone, a PLK4 inhibitor, to determine centriole assembly in S phase in the PLK4-overexpressing cells and the triple KO cells (Wong et al., 2015). As expected, the number of centrioles in control cells remained two at S phase (Figure 3E). The average number of centrioles in PLK4-overexpressing cells started with seven and increased to ten during S phase (Figure 3E). However, the S phase assembly of the centrioles was inhibited with the centrinone treatment (Figure 3E). The average number of centrioles in the triple KO cells were 3.5 and this number hardly increased even at S phase, (Figure 3E). Centrinone had a little effect on the triple deletion cells, confirming that the triple KO cells hardly assemble procentrioles in S phase (Figure 3E).

### Centriole-to-centrosome conversion in the precociously separated centrioles

We investigated centriole-to-centrosome conversion after precocious centriole separation at M phase. When *PCNT*-deleted cells were arrested at prometaphase with STLC, their centrioles readily separated (Figure 2D; Kim et al., 2019). We determined localization of CEP295 and CEP152 in the precociously separated centrioles at M phase. The results showed that about halves of the *TP53;PCNT*-deleted and *TP53;PCNT;CEP215*-deleted cells included three and more CEP295 and CEP152 signals in their centrioles (Figure 4). On the other hand, CEP295 and CEP152 signals were detected at a centriole pair in most of the *TP53*- and *TP53;CEP215*-deleted cells (Figure 4). These results suggest that daughter centrioles readily convert to centrosomes even at M phase as soon as they are separated from the mother centrioles.

**Figure 4.**
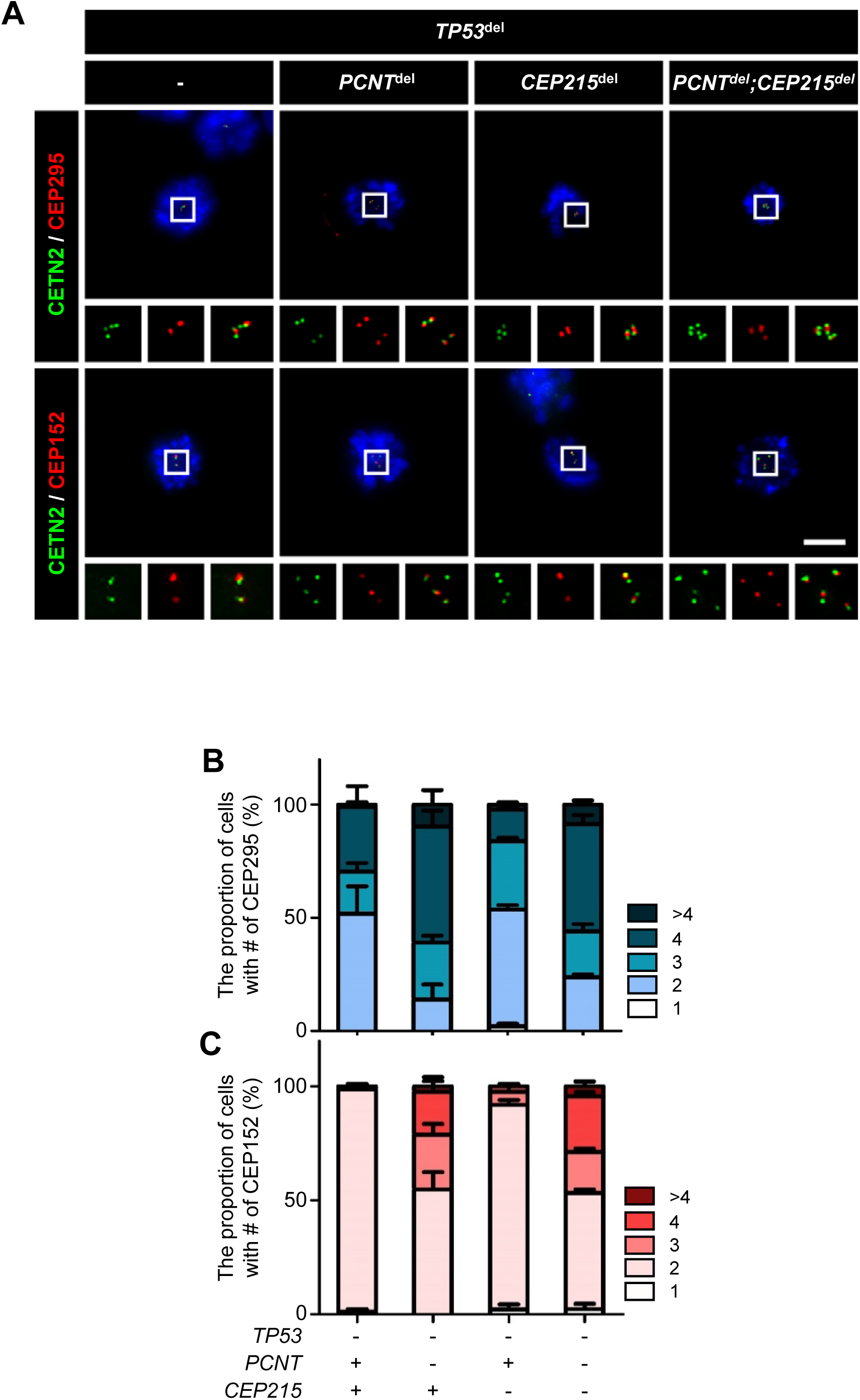
Centriole-to-centrosome conversion in precociously separated centrioles during M phase. (A) The triple KO cells were treated with STLC for 10 h and coimmunostained with CETN-2 (green) and CEP295 or CEP152 (red). Nuclei were stained with DAPI (blue). Scale bar, 10 μm. (B) The numbers of CEP295 and CEP152 signals were counted in the cells. Greater than 30 cells per group were analyzed in three independent experiments. Values are means and SEM.

### Defective centriole-to-centrosome conversion in the triple KO cells

Presence of only two CEP152-positive centrioles out of multiple ones in the triple KO cells suggests that only a pair of centrioles are able to recruit PLK4 for generation of new procentrioles during S phase. In order to examine intactness of the centrioles in the triple KO cells, we performed coimmunostaining analysis with CEP152 and selected centrosome proteins (Figure 5A). As expected, most of centrioles in the control cells were CEP152-positive and also coimmunostained with antibodies specific to CEP295, CEP192, CEP135 and γ-tubulin (Figure 5B). Most of the multiple centrioles in the PLK4-overexpressing cells were immunostained with all the antibodies we used (Figure 5B). However, only two centrioles in the triple KO cells were positive to CEP152 (Figure 5B). Furthermore, CEP295, CEP192, CEP135 and γ-tubulin were detected almost exclusively at the CEP152-positive centrioles (Figure 5B). These results strongly suggest that only a pair of the CEP152-positive centrioles are intact mother centrioles whereas the rest of them are defective centrioles in the triple KO cells.

**Figure 5.**
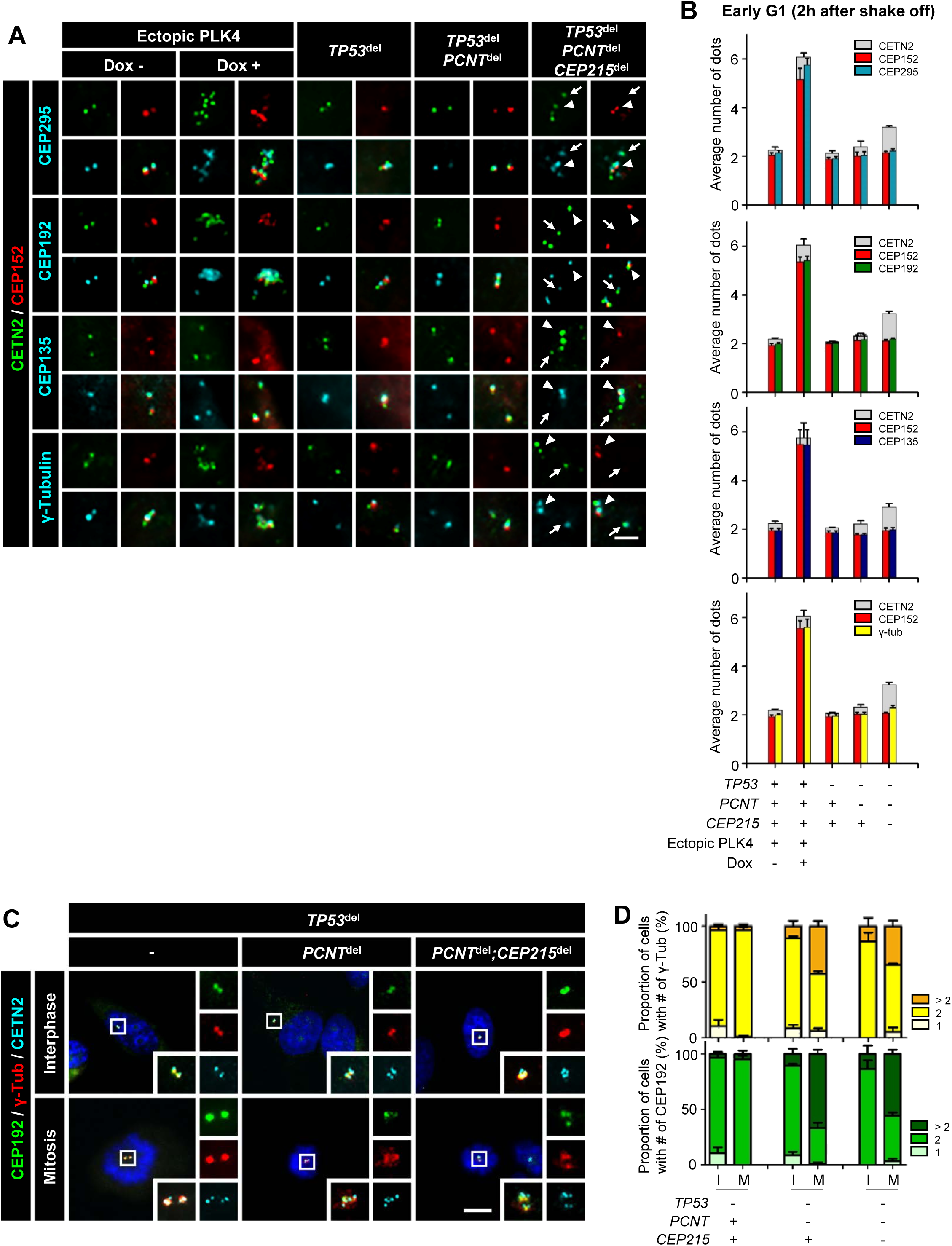
Determination of intact centrioles in the triple KO cells. (A) The PLK4-overexpressing and triple KO cells at G1 phase were coimmunostained with antibodies specific to CEP295, CEP192, CEP135 and γ-tubulin (cyan), along with CETN2 (green) and CEP152 (red). Scale bar, 2 μm. Arrows and arrowheads mark the CEP152-positive and -negative centrioles, respectively. (B) The number of centrioles with CEP152 and the indicated antibodies were counted. (C) The KO cells at interphase and mitosis were coimmunostained with antibodies specific to CEP192 (green), γ-tubulin (red) and CETN2 (cyan). Nuclei were stained with DAPI (blue). Scale bar, 10 μm. (D) The number of CEP192 and γ-tubulin signals were counted in cells at interphase (I) and mitosis (M). Greater than 30 cells per group were analyzed in three independent experiments. Values are means and SEM.

We performed a similar coimmunostaining analysis with the KO cells at mitosis (Figure 5C, D). Consistent with the previous results, the control as well as the KO cells at interphase had two centrosomes positive to CEP192 and γ-tubulin (Figure 5C, D). In the mitotic control cells, both CEP192 and γ-tubulin signals were detected at two pairs of centrioles (Figure 5C, D). The double and triple KO cells had centrioles separated and frequently amplified at mitosis. Both the CEP192 and γ-tubulin signals were detected in many of the separated and amplified centrioles of the deletion cells at mitosis (Figure 5D). These results suggest that separated centrioles in the triple KO cells may have an ability to organize microtubules during mitosis.

### Defective microtubule organization in the triple KO cells

We performed microtubule regrowth assays to determine biological activities of the centrosomes in the PLK4-overexpressing cells and the triple KO cells. In control cells, microtubules started to be organized from both centrosomes present at G1 phase (Figure 6A, B). About 6 centrosomes are present in the PLK4-overexpressing cells and 91% of them were able to organize microtubules (Figure 6A, B). However, in the tripled KO cells, only 73% of the centrosomes organized microtubules, leaving 27% of them without an activity (Figure 6A, B). These results suggest that a significant fraction of the centrosomes in the triple KO cells have functional defects in microtubule organization during interphase.

**Figure 6.**
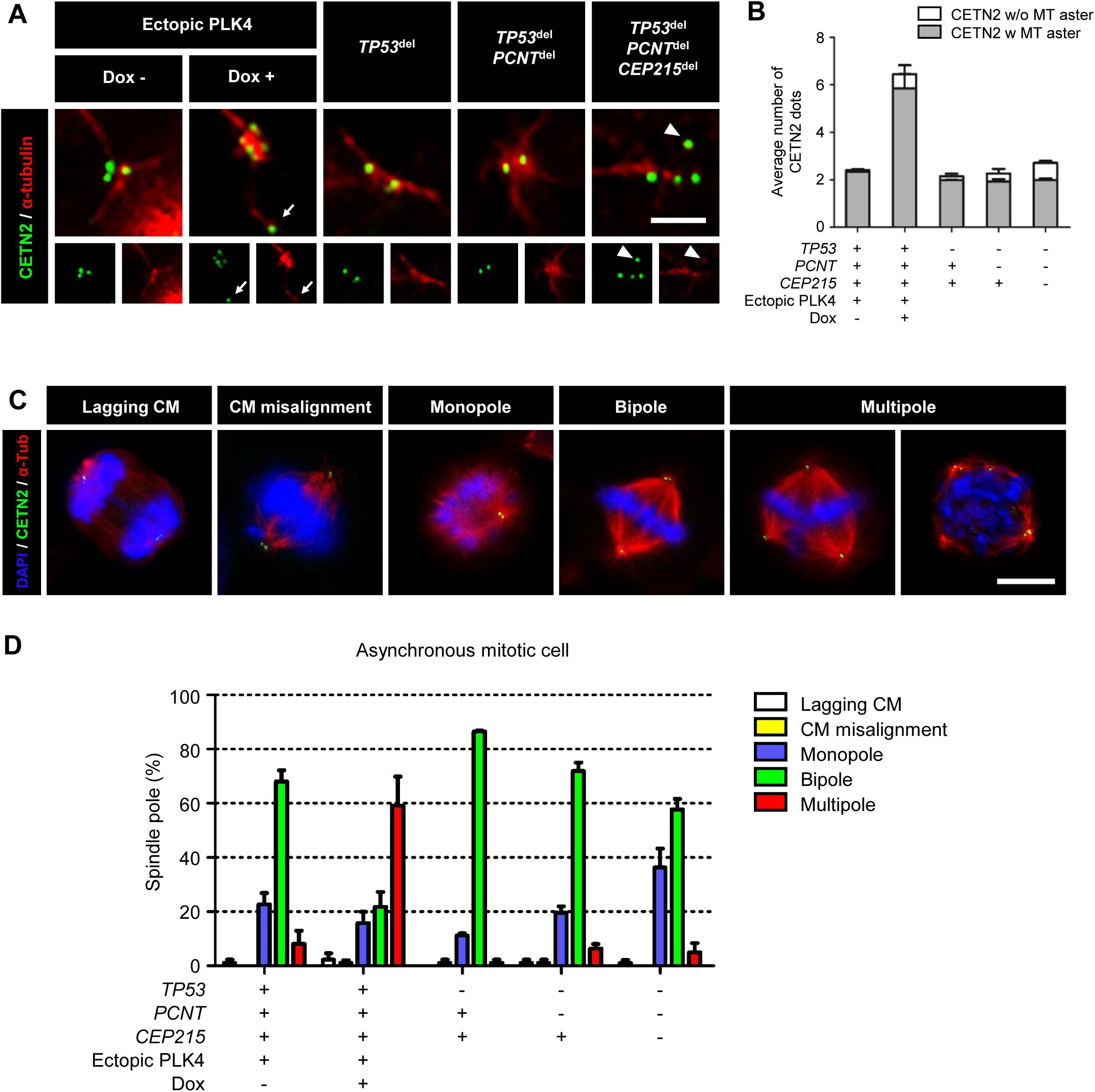
MTOC activities in the supernumerary centrioles. (A) The PLK4-overexpressing and triple KO cells at G1 phase were subjected to microtubule regrowth assays. The cells were coimmunostained with antibodies specific to CETN2 (green) and α-tubulin (red). Scale bar, 2 μm. (B) The number of CETN2 dots with microtubule asters are counted. (C) The PLK4-overexpressing and triple KO cells at mitosis were subjected to coimmunostaining analysis with antibodies specific to CETN2 (green), α-tubulin (red) and DAPI (blue). Representative abnormalities of the spindle poles were shown. Scale bar, 10 μm. (D) Mitotic cells with abnormal spindle poles were counted. (B, D) Greater than 30 cells per group were analyzed in three independent experiments. Values are means and SEM.

We determined spindle configurations in mitotic cells of the PLK4-overexpressing and triple KO cells. Spindle pole phenotypes are categorized into five groups following Watanabe et al. (2019); lagging chromosome, chromosome misalignment, monopole, bipole and multipole (Figure 6C). As expected, most of the mitotic control cells formed bipolar spindles (Figure 6D). In PLK4-overexpressing cells, about 60% of mitotic cells formed multipoles (Figure 6D). On the other hand, the proportion of monopoles were significantly increased up to 40% in the triple KO cells but the mitotic cells with multipoles were insignificant (Figure 6D). These results suggest that many of the separated centrioles in the triple KO cells have limited ability to function as spindle poles during mitosis.

## DISCUSSION

In this work, we generated *TP53, PCNT* and *CEP215* triple KO cell lines and determined their phenotypes at the centrosome. We observed that centrioles in the triple KO cells precociously separated and amplified at M phase. Many of the triple KO cells maintained supernumerary centrioles throughout the cell cycle. However, the number of centrioles did not double during S phase. It is likely that, in the triple KO cells, supernumerary centrioles, many of which are assembled during M phase, lack an ability to function as templates for centriole assembly during S phase (Figure 7).

**Figure 7.**
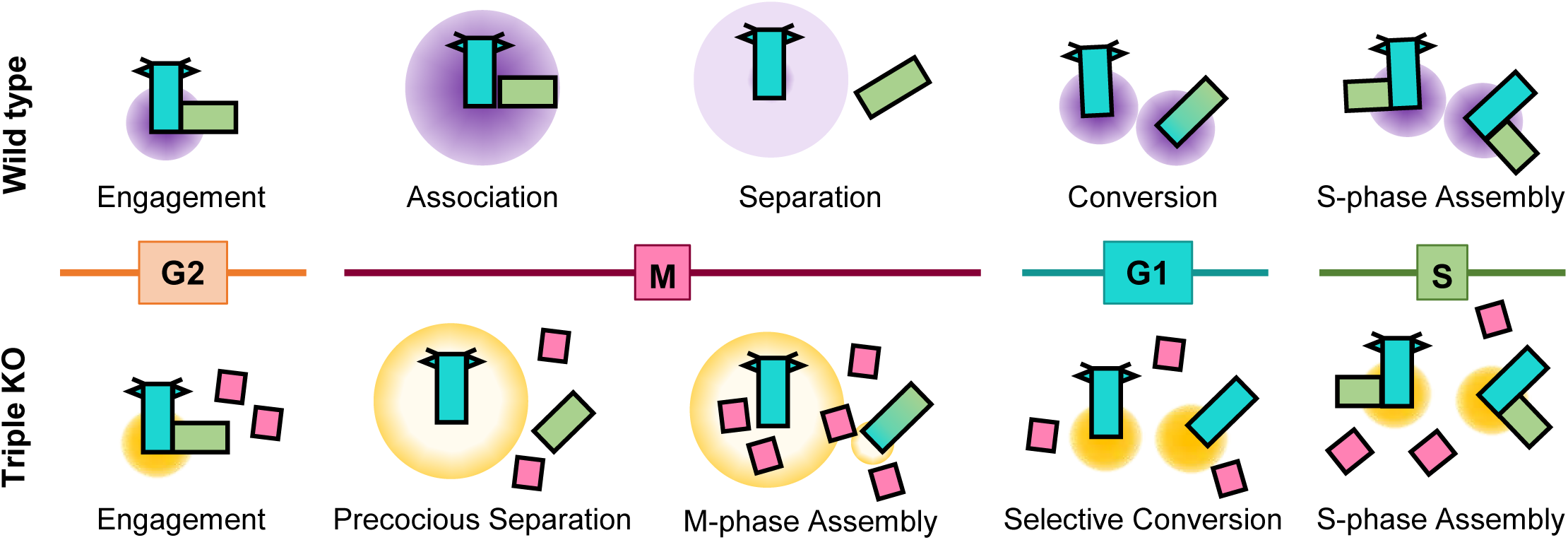
Model. At early M phase, daughter centrioles readily disengage from mother centrioles, but remain associated surrounded by mitotic PCM. Daughter centrioles eventually separate from the mother centrioles after PCM is disintegrated at the end of mitosis. Deletion of *PCNT* and *CEP215* makes mitotic PCM disorganized. As results, daughter centrioles precociously separate from mother centrioles at M phase and centriole amplification occurs. The M-phase-assembled centrioles, however, cannot convert to mother centrioles during mitotic exit. They do not organize microtubules, nor function as templates for nascent centriole assembly in subsequent S phase, and are detected throughout the cell cycle.

Since PCNT and CEP215 are major PCM components in the human centrosome, deletion of them might result in disorganized structure of mitotic PCM. As previously shown, most of the *PCNT*-deleted cells revealed precocious centriole separation during mitosis and a small fraction of them have multiple centrioles (Kim et al., 2019). Centriole-to-centrosome conversion readily follows after precocious centriole separation at M phase (Tsuchiya et al., 2016). However, less than a half of *CEP215*-deleted cells had precocious centriole separation without amplification, suggesting that integrity of the mitotic PCM is more or less maintained in the absence of CEP215. In fact, we and others reported that mitotic centrosomes are relatively intact in the CEP215-depleted cells, except that their centrioles are frequently displaced from spindle poles during mitosis (Lee and Rhee, 2010; Barr et al., 2010; Chavali et al., 2016; Gambarotto et al., 2019). Nonetheless, deletion of *CEP215* enforced the *PCNT* KO phenotypes, suggesting that PCNT and CEP215 cooperatively constitute mitotic PCM to maintain centriole association during M phase (Buchman et al., 2010; Kim and Rhee, 2014). Monopolar spindles in the triple KO cells should be attributed to a low microtubule organizing activity, due to a limited accumulation of mitotic PCM (Lee and Rhee, 2010; Figure 6d).

The M-phase-assembled centrioles in the triple KO cells failed to duplicate during S phase. In contrast, the S-phase-assembled centrioles in PLK4-overexpressing cells doubled in the next S phase. These results strongly suggest that majority of the M-phase-assembled centrioles lack an ability to function as template for nascent centriole assembly during S phase. In fact, only two out of multiple centrioles in the triple KO cells are positive to CEP152 which is a scaffold for PLK4 in mother centrioles. Known preceding components for the centriole-to-centrosome conversion, such as CEP135, CEP295 and CEP192, were also detected almost exclusively at the CEP152-positive centrioles, indicating that only a single out of many centrioles in a triple KO cell is able to convert from centriole to centrosome during mitotic exit. We suspect that two CEP152-positive centrioles may be generated at the previous S phase, while the other centrioles are assembled at M phase. It remains to be investigated why the M-phase-assembled centrioles cannot be converted to centrosome during mitotic exit. A possibility may be that centriole assembly processes cannot be compressively proceeded within a short M phase. As a result, the M-phase assembled centrioles may not be able to recruit a series of centrosomal proteins necessary for the conversion until the end of mitosis. It remains to be identified what factors are critically absent for the M-phase-assembled centrioles to undergo conversion.

Once a daughter centriole converts to a mother centriole during mitotic exit, it acquires an ability to recruit PCM (Wang et al., 2011; Fu et al., 2016). We observed that almost all amplified centrioles in the PLK4-overexpressing cells can organize microtubules. As a result, majority of the PLK4-overexpressing cells formed multipoles in M phase. On the other hand, a significant proportion of centrioles in the triple KO cells failed to organize microtubules in interphase. These results also support the notion that the M-phase-assembled centrioles cannot convert to centrosomes during mitotic exit. Consequently, they hardly function as microtubule organizing centers during the cell cycle. Nonetheless, we do not rule out the possibility that a fraction of the M-phase-assembled centrioles may acquire an ability to organize microtubules especially during mitosis. In fact, we found the γ-tubulin signals in some of the extra centrioles of the triple KO cells at M phase.

Supernumerary centrioles are common in cancer cells (Chan, 2011; Marteil et al., 2018). It is not clear whether supernumerary centrosomes are sufficient to drive tumor development or not (Vitre et al., 2015). It remains to be investigated how centrioles are amplified and how supernumerary centrioles are maintained in tumor cells (Kwon et al., 2013). Supernumerary centrioles may be generated by a number of non-physiological pathways, in addition to PLK4 overexpression. In this work, we revealed that M-phase-assembled centrioles are generated in cells which lack critical PCM proteins. The supernumerary centrioles which are generated during M phase may be dormant with little biological activity, and, therefore, are maintained throughout the cell cycle. However, since these centrioles can also recruit γ-tubulin at M phase, they have a chance to cause mitotic defects and aneuploidy.

## MATERIALS AND METHODS

### Cell culture, generation of deleted cell lines and synchronization

The deleted cell lines were made in the Flp-In T-Rex Hela cells (Kim et al., 2019). *CEP215* was deleted using CRISPR/Cas9 technique in the *PCNT, TP53*-double deleted cells expressing DD-PCNT (Kim et al., 2019). The gRNA sequences for *CEP215* deletion are (5’-ccagggacggtgacgtcctcttc-3’) and (5’-ctgcagccgctgagcgtcccagg-3’). HeLa cells were cultured in DMEM (LM001-05; Welgene, Gyeongsangbuk-do, South Korea) with 10% FBS (S101-01; Welgene). For mitotic synchronization, cells were sequentially treated with 2mM thymidine (T9250; Sigma-Aldrich, St. Louis, MO) and 5μM STLC (2191; Tocris, Bristol, United Kingdom). For the time course experiment, cells were treated with doxycycline for 24 hours after seeding to induce ectopic PLK4 expression, washed out and cultured for another 24 hours. Mitotic cells were obtained with a gentle shake-off from asynchronous cell plates and collected with a warm medium at indicated time points.

### Microtubule regrowth assay

Mitotic cells were obtained with mitotic shake-off and cultured for 2 h to reach at early G1 phase. The cells were treated with 5μM of nocodazole (M1404; Sigma-Aldrich) for 2 hours at 37°C, placed on ice for 1 hour, and then transferred to a warm medium for microtubule growth. The cells were fixed with PEM buffer (80mM PIPES pH6.9, 1mM MgCl2, 5mM EGTA, 0.5% Triton X-100) for 10 minutes at room temperature, incubated in phosphate-balanced buffer with 0.5% Triton-X (PBST) for 5 minutes to increase permeability, and subjected to immunostaining with antibodies specific to α-tubulin and centrin-2.

### Antibodies

The antibodies specific to centrin-2 [immunocytochemistry (ICC) 1:1000; 04-1624; Merck Millipore, Billerica, MA], CEP295 (ICC 1:500; 122490; Abcam, Cambridge, MA), CEP192 [ICC 1:1000, immunoblot (IB) 1:500; A302-324A; Bethyl Laboratories, Montgomery, TX], CEP152 (ICC 1:500, IB 1:100; 183911; Abcam), GAPDH (IB 1: 20,000; AM4300; Life Technologies, Carlsbad, CA), SAS-6 (ICC 1:200, IB 1:100; sc-376836; Santa Cruz Biotechnology, Dallas, TX), α-tubulin (ICC 1:2000, IB 1: 20,000; ab18251; Abcam), γ-tubulin (ICC 1:1000, IB 1:2000; 11316; Abcam) were purchased. The antibodies specific to CEP215 (Lee and Rhee, 2010; ICC 1:2000, IB 1:500), PCNT (Kim and Rhee, 2011; ICC 1:2000, IB 1:2000), CEP135 (Kim et al., 2008; ICC 1:2000) and CPAP (Chang et al., 2010; ICC 1:100; IB 1:500) were previously described. Secondary antibodies conjugated with fluorescent dyes (ICC 1:1000; Alexa Fluor 488, 594 and 647; Life Technologies) and with horseradish peroxidase (IB 1:10,000; Sigma-Aldrich or Millipore) were purchased.

### Immunostaining analysis

Cells on a cover glass (0117520; Paul Marienfeld, Lauda-Königshofen, Germany) were fixed with cold methanol for 10 min at 4°C, washed with cold PBS, and blocked with blocking solution (3% bovine serum albumin, and 0.3% Triton X-100 in PBS) for 30 min. The samples were incubated with primary antibodies for 1 h, washed with 0.1% PBST, incubated with secondary antibodies for 30 minutes, washed, and treated with 4,6-diamidino-2-phenylindole (DAPI) solution for up to 2 minutes. The cover glasses were mounted on a slide glass with ProLong Gold antifade reagent (P36930; Life Technologies). Images were observed with fluorescence microscopes with a digital camera (Olympus IX51) equipped with QImaging QICAM Fast 1394 and processed in ImagePro 5.0 (Media Cybernetics). ImagePro 5.0 (Media Cybernetics), Photoshop CC (Adobe) and ImageJ 1.51k (National Institutes of Health) were used for image processing. For measuring fluorescence intensities at centrosome, all images were obtained in an identical setting with the same exposure time. ImageJ was used for measuring and the background signals were subtracted from the centrosomal signals.

### Live cell imaging

CQ1 benchtop high-content analysis system was used for imaging live cells. The *CETN2-Dendra2* plasmid which had been kindly provided by Alwin Krämer (Löffler et al., 2012) were stably transfected into the cells. The cells were synchronized at M phase with sequential treatments of thymidine and STLC, activated with 405nm wavelength for 10 sec and recorded with 488nm and 561nm wavelengths for up to 2 h.

### Immunoblot analysis

Cells were washed with PBS, lysed on ice for 10 min with RIPA buffer (1% Triton X-100, 150 mM NaCl, 0.5% sodium deoxycolate, 0.1% SDS, 50 mM Tris-HCl at pH 8.0, 1 mM Na3VO4, 10 mM NaF, 1 mM EDTA and 1 mM EGTA) containing a protease inhibitor cocktail (Sigma-Aldrich, P8340) and centrifuged for 10 minutes at 4°C. A fraction of the supernatants was used for the Bradford assays, and the rest were mixed with 4×SDS sample buffer (250 mM Tris-HCl at pH 6.8, 8% SDS, 40% glycerol and 0.04% bromophenol blue) and 10 mM DTT (0281-25G; Amresco). The mixtures were boiled for 5 min. For PCNT, CEP215, and CEP152, 3% stacking gel and 4% separating gel were used with 20mg of protein samples. For SAS-6 and CPAP, 5% stacking gel and 8% separating gel were used with 20mg of protein samples. The rest are loaded in 5% stacking gel and 10% separating gel with 10mg of protein samples. Proteins at gels were transferred to Protran BA85 nitrocellulose membranes (10401196; GE Healthcare Life Sciences). The membranes were blocked with a blocking solution (5% nonfat milk in 0.1% Tween 20 in TBS or 5% bovine serum albumin in 0.1% Tween 20 in TBS) for 2 h, incubated with primary antibodies diluted in blocking solution for overnight at 4°C, washed with TBST (0.1% Tween 20 in TBS), incubated with secondary antibodies in blocking solution for 30 min and washed again. ECL reagent (LF-QC0101; ABfrontier, Seoul, South Korea) and X-ray films (CP-BU NEW; Agfa, Mortsel, Belgium) were used to detect the signals.

---

**Figure 1 Supplement Figure 1.**
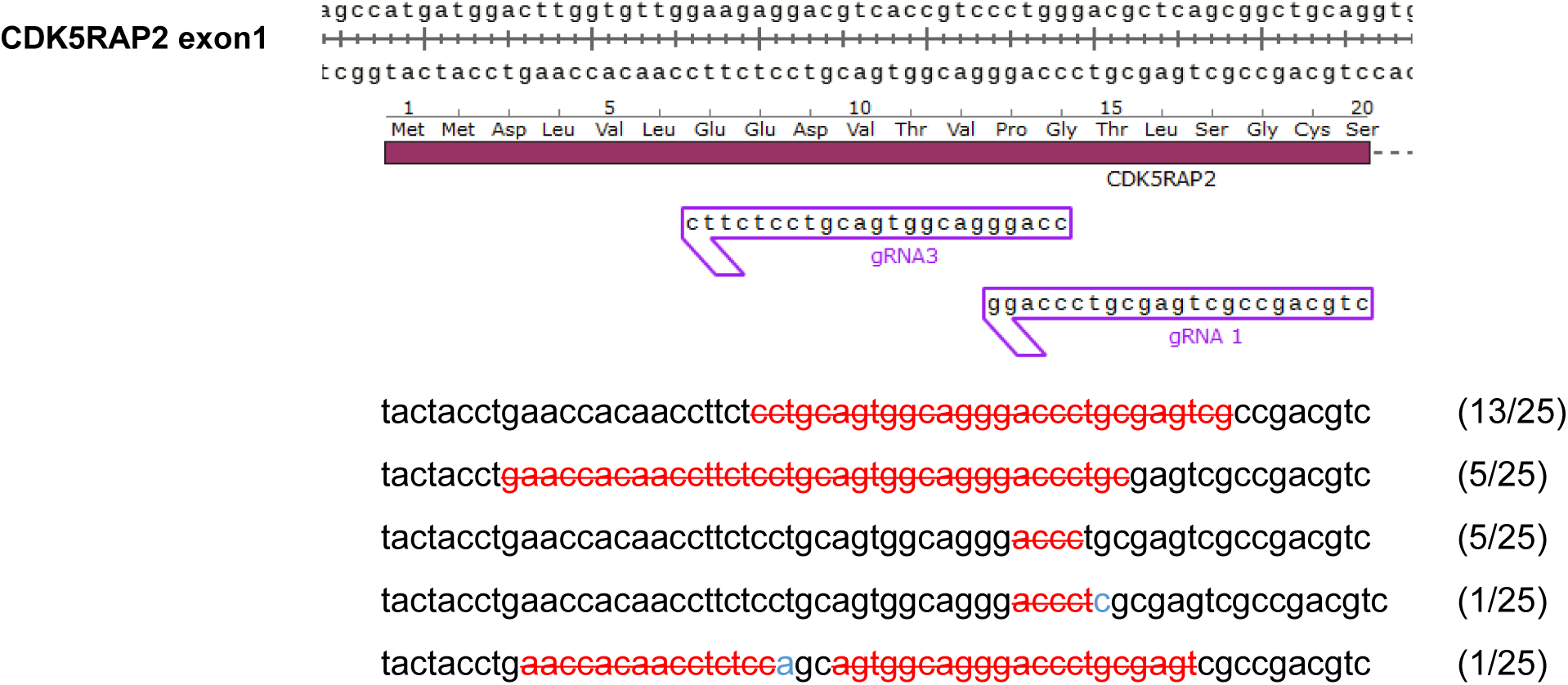
Deletion of *CEP215* in HeLa cells. The *CEP215* gene was deleted using the CRISPR/CAS9 technique. Guide RNAs and indel types were indicated. Insertions and deletions were marked with blue and red colors, respectively.

**Figure 2 Supplement Movie 1.**
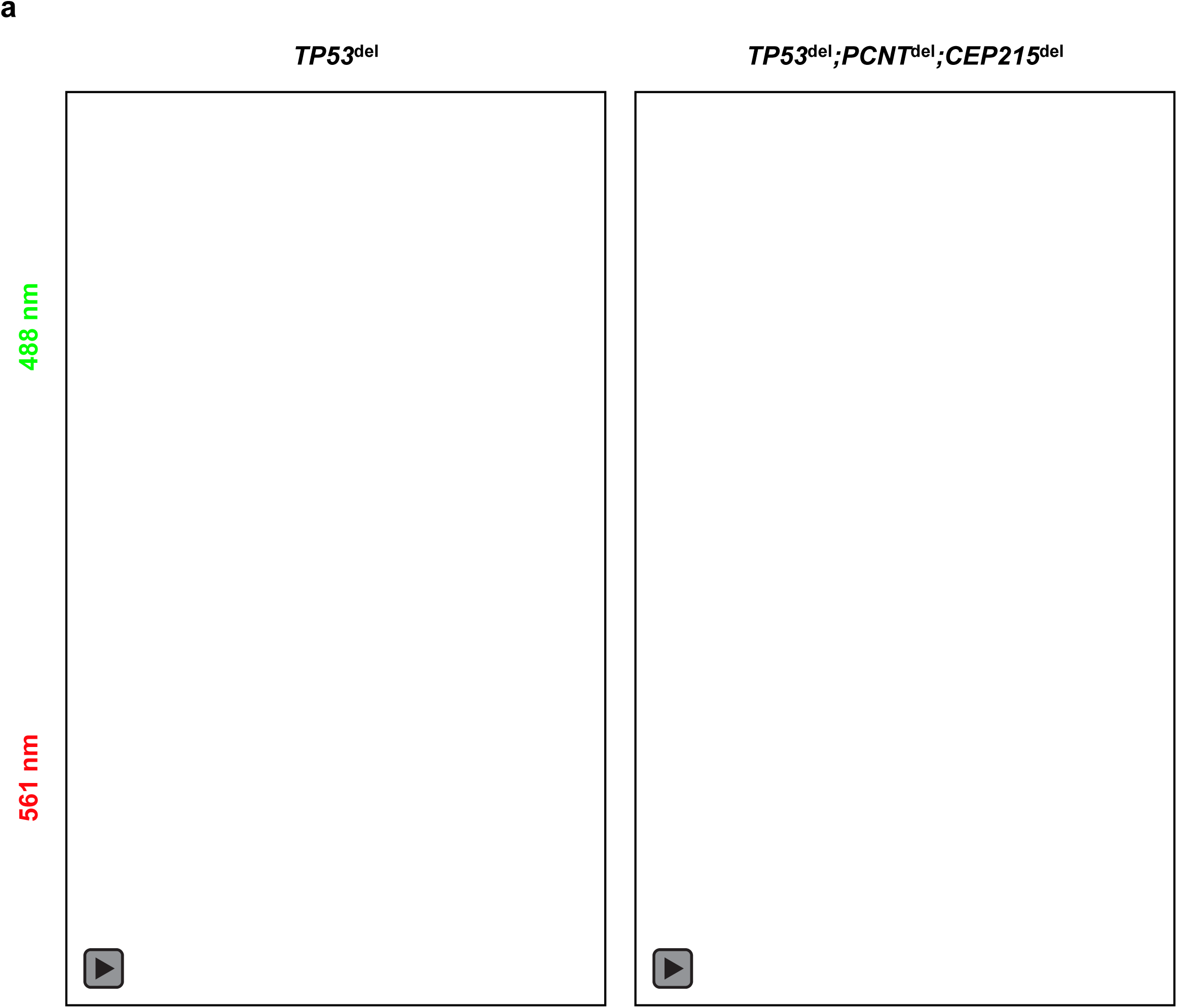
Live cell imaging of CETN2-Dendra2 in the triple KO cells. The CETN2-Dendra2-expressing triple KO cells were treated with thymidine for 24 h followed by STLC for 8 h. After a light activation, the CETN2-Dendra2 signals were recorded in 15 min-intervals for up to 2 h. Preexisting centrioles were detected with both 488 nm and 561 nm wavelengths, while de novo centrioles (arrow) were detected only with 488 nm wavelength.

